# The enteroviral protease target LSM14A operates outside of P-bodies to augment antiviral innate immunity

**DOI:** 10.64898/2026.05.11.723630

**Authors:** Alan Wacquiez, Hicham Hboub, Da-Yuan Chen, Alexander H. Tavares, Chia-Ming Su, Jocelyn De Paz, Marc Semaan, Zhen Ding, Joseph Zaia, Manveen K. Sethi, Shawn M. Lyons, Mohsan Saeed

## Abstract

Antiviral innate immune networks in human cells comprise core components that serve as central signaling hubs and several context-dependent modulators whose role may be virus- or tissue-specific. One such modulator is LSM14A, which potentiates innate immune response but is not essential. We recently showed that enteroviruses deploy their protease activity to cleave LSM14A, thereby disabling its antiviral function. In this study, we probe the molecular mechanism by which LSM14A contributes to innate immunity. We show that although LSM14A predominantly localizes to processing bodies (P-bodies; PBs), this localization is not essential for its innate immune function. Likewise, association with peroxisomes does not contribute to its immune activity. Instead, an unbiased systems-level interactomic analysis reveals a distinct cohort of LSM14A-associated proteins that assemble outside canonical PBs and peroxisomes following infection with Sendai virus, a robust inducer of innate immunity. Functional interrogation of these interactors demonstrate that several are essential for LSM14A-dependent amplification of antiviral signaling. Together, these findings uncover a functional axis of LSM14A that operates independently of its canonical subcellular localizations and is mediated through a specialized interaction network, improving our understanding of how this protein reinforces the antiviral innate immune system.

## INTRODUCTION

Antiviral innate immune networks comprise an array of cellular proteins with specialized functions. While some of these proteins represent well-defined core sensors, adaptors, and effectors that constitute the central hubs of innate immune signaling^1,2^, others operate at the periphery to fine-tune antiviral responses. These so-called auxiliary proteins do not typically initiate signaling on their own but modulate the strength, timing, and spatial organization of canonical pathways in a context- or cell type-dependent manner. They may act by scaffolding transient protein-protein interactions^3–5^, regulating RNA or protein stability^6–8^, or organizing signaling complexes within specialized subcellular microenvironments, such as BPs^9,10^, stress granules (SGs)^11,12^, or organelle contact sites^13–15^. Additionally, their contribution can vary with the virus or the cell type used, enabling tailored responses to distinct pathogenic challenges in specific cell states. As a result, these peripheral regulators can markedly enhance innate immune outputs, providing an additional layer of control that expands the flexibility and specificity of host antiviral defenses.

Viruses have evolved strategies to target both the core components and peripheral regulators of antiviral defense systems. We recently employed a systems-level approach to demonstrate that enteroviruses use their proteases to target a number of innate immune-related proteins during infection^16^. One such protein is LSM14A, which is a critical PB component^17–19^. Our functional assays showed that while the full-length LSM14A enhances immune responses to viral infection, its N- and C-terminal cleavage products, generated by the enteroviral 2A protease (2A^pro^), fail to perform this function^16^, suggesting that the viral protease activity blocks LSM14A’s innate immune activity. However, despite studies clearly showing the role of LSM14A in antiviral immunity^10,20–22^, and viruses targeting this protein to blunt the innate immune pathway^16,23^, the mechanisms by which LSM14A reinforces antiviral immune responses remain poorly understood.

Li *et al.* reported that LSM14A translocates to peroxisomes following viral infection, where it is proposed to interact with RIG-I, MAVS, and STING to enhance innate antiviral signaling^10^. However, whether this relocalization is required for LSM14A’s immune-enhancing activity remains untested. In another study, LSM14A^-/-^ mouse dendritic cells were used to demonstrate that LSM14A regulates STING protein levels in a tissue- and cell type-specific manner, apparently by modulating STING mRNA processing; LSM14A deficiency did not affect the abundance of STING pre-mRNA, but reduced mature mRNA levels, indicating a defect in mRNA maturation^20^. However, key mechanistic questions remain unresolved: how LSM14A, which is primarily localized in the cytoplasm, controls STING mRNA processing, what underlies its cell type-specific effects, and whether this activity is sufficient to explain LSM14A-mediated amplification of innate immune responses, particularly in the context of RNA virus infection.

LSM14A was originally identified in salamander eggs as a ∼55 kDa RNA-associated protein—termed RAP55—in messenger ribonucleoprotein (mRNP) complexes that harbor translationally repressed mRNAs^24^. These mRNAs are mobilized at defined stages of embryonic development, with RAP55 contributing to their repression within mRNP complexes^25^. Subsequent studies in human cells revealed that the protein, renamed LSM14A due to the presence of an N-terminal LSM domain^26^, resides within PBs and plays essential roles in the formation of these granules by interacting with two key PB constituents, the eIF4E-binding protein 4E-T and the DEAD-box helicase DDX6^27^. Structural analyses have defined the molecular interface between these proteins: the LSM domain of LSM14A is wrapped by a bi-partite motif in the C-terminus of 4E-T, while LSM14A engages DDX6 mainly through their respective C-terminal regions^27^. In its unbound state, LSM14A is largely intrinsically disordered, except for the N-terminal LSM domain, a central phenylalanine-aspartate-phenylalanine (FDF) motif, and a C-terminal RGG motif^28^. Two additional highly conserved, yet functionally uncharacterized motifs, FFD and TFG, have been identified between the FDF and RGG motifs^26^.

We previously showed that LSM14A overexpression amplifies Sendai virus (SeV)-induced innate immune responses^16^. This effect operates through the canonical innate immune pathway, as depletion of RIG-I, the primary sensor of SeV RNA, and the adaptor protein MAVS abrogated LSM14A-mediated immune amplification. In contrast, knockdown of MDA5 had no effect. Notably, genetic depletion of LSM14A did not impair SeV-induced innate immune activation, indicating that LSM14A is not essential for pathway activation but functions as a potent enhancer of the response.

In this study, we establish new tools demonstrating that LSM14A exerts antiviral activity against enteroviruses. We also demonstrate that LSM14A’s immune-enhancing function is independent of canonical PBs and peroxisomes. Systematic mapping of LSM14A interaction partners outside these compartments identified several functionally relevant proteins whose depletion impairs LSM14A-mediated immune amplification. Collectively, these findings provide mechanistic insights into how LSM14A potentiates innate immune signaling.

## RESULTS

### LSM14A possesses antiviral activity

We previously showed that LSM14A is targeted by the enteroviral 2A^pro^ for proteolytic cleavage^16^, suggesting that LSM14A may function as an antiviral factor against enteroviruses. However, overexpression of LSM14A did not inhibit viral infection. We attributed this lack of effect to the concurrent cleavage of other innate immune components, particularly MAVS, during infection. This hypothesis was reinforced by the observation that LSM14A fails to amplify the innate immune response in MAVS knockout cells^16^. To overcome this challenge, we generated recombinant poliovirus (PV) expressing catalytically inactive 2A^pro^ (PV^2A-mut^), using a recently described approach^29^. As the proteolytic activity of 2A^pro^ is essential for processing at the VP1-2A^pro^ junction, we replaced the 2A^pro^ cleavage site at this junction with the sequence corresponding to the 3CD cleavage site recognized by 3C protease (3C^pro^).

To test the inhibitory effect of LSM14A against enteroviruses, we generated human intestinal epithelial Caco-2 cells stably expressing LSM14A and infected them with PV^2A-mut^. LSM14A overexpression reduced viral titer by ∼5-fold compared to control cells expressing the empty vector (Fig. 1A). Overexpression of MAVS, used as a positive control, suppressed viral replication by 10-fold. Together, these results indicate that LSM14A is an antiviral factor, and enteroviruses have evolved to deploy their 2A^pro^ to inactivate this protein.

**Fig. 1.**
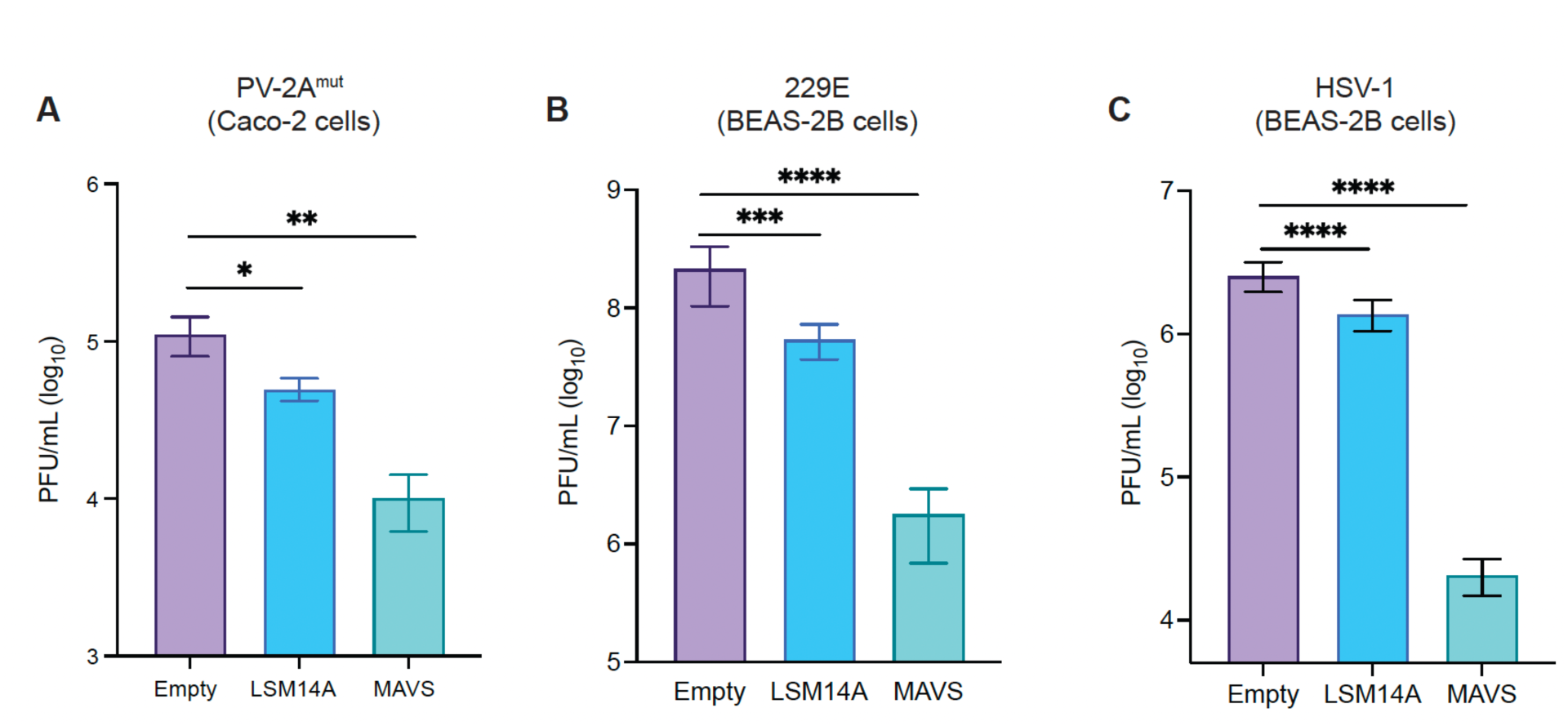
LSM14A overexpression suppresses virus infection. **A.** Caco-2 cells stably expressing empty pLOC, pLOC-LSM14A or pLOC-MAVS were infected with PV-2A^mut^ at an m.o.i of 0.01, and the viral titer in the culture medium was measured at 24 h.p.i. **B, C.** BEAS-2B cells stably expressing empty pLOC, pLOC-LSM14A or pLOC-MAVS were infected with human coronavirus 229E (B) or with HSV-1 (C) at an m.o.i of 0.01 followed by virus titration in the culture medium at 24 h.p.i. Statistical significance was determined using two-tailed non-parametric *t*-test. *P < 0.05, **P < 0.001, ***P < 0.0001.

Given that LSM14A enhances innate immune response to viral infection, we hypothesized that it would exert broad antiviral activity. To test this, we used human airway epithelial BEAS-2B cells which mount a robust innate immune response upon virus infection^30^. LSM14A overexpression in these cells significantly reduced replication of the RNA virus human coronavirus 229E (Fig. 1B) and the DNA virus herpes simplex virus 1 (HSV-1) (Fig. 1C). Together, these findings establish LSM14A as a potent antiviral factor with activity across viruses with distinct genome types.

### PBs or peroxisomes are not required for LSM14A’s innate immune function

LSM14A localizes to PBs, and together with DDX6 and 4E-T, constitutes a core component of these structures: depletion of any of these proteins disrupts PB formation^27,31^. Immunofluorescence analysis of 293T cells using anti-LSM14A antibodies revealed distinct punctate structures characteristic of PBs, and the abundance of these puncta markedly increased upon treatment with sodium arsenite, a known inducer of PB formation (Fig. 2A). These puncta co-stained with DDX-6 and 4E-T, confirming their *PB* identity (Fig. 2B).

**Fig. 2.**
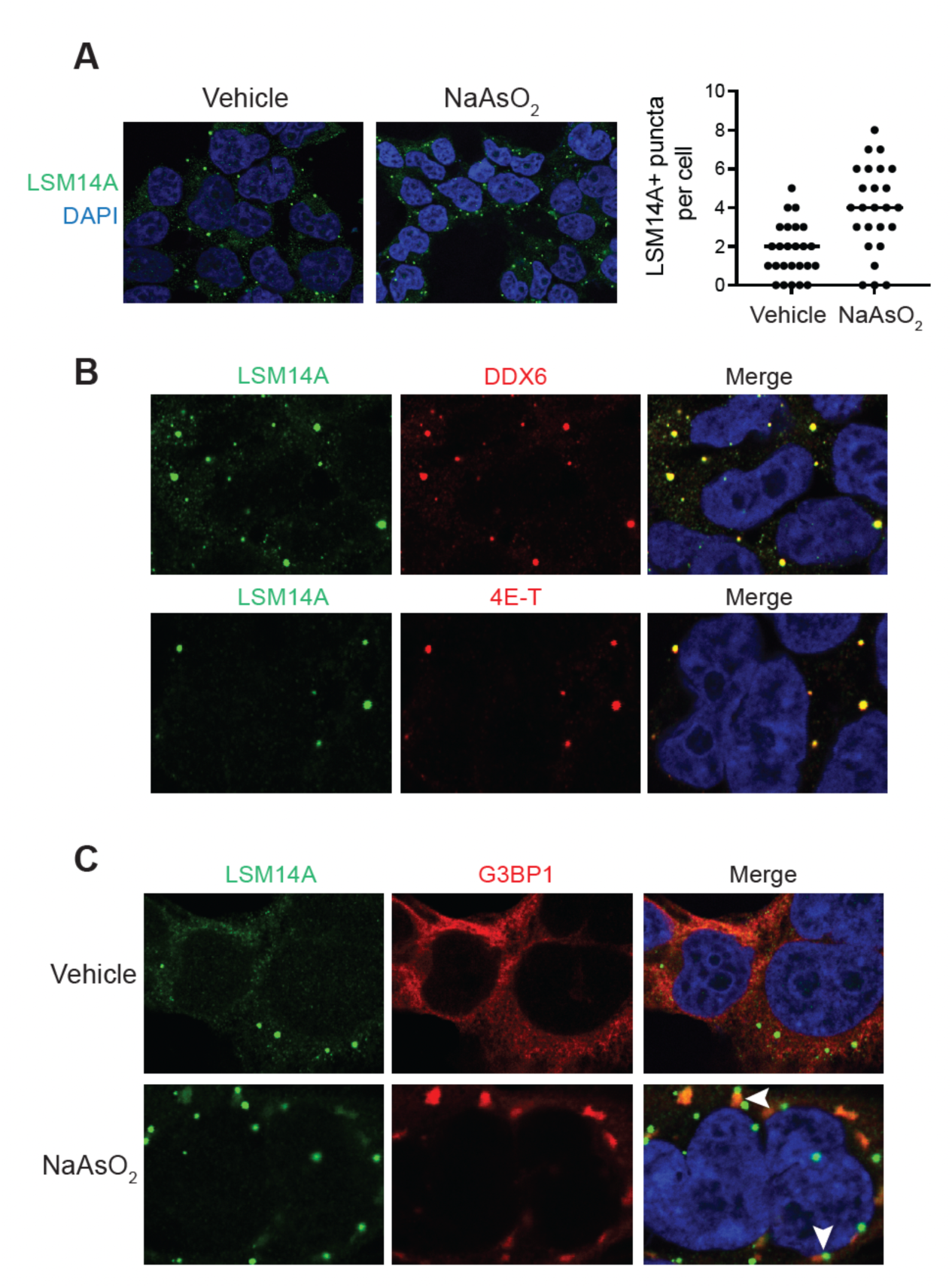
LSM14A localizes to PBs. **A.** 293T cells were either treated with sodium arsenite (NaAsO_2_) (100 µM for 1h at 37°C) or DMSO (vehicle), followed by staining with anti-LSM14A antibody. The graph on the right shows the number of LSM14A+ puncta in 25 randomly selected cells. Each dot represents a cell. Statistical significance was determined using two-tailed non-parametric *t*-test. *P < 0.05. **B.** 293T cells were stained with anti-LSM14A and either anti-DDX6 antibodies (Top panel) or with anti-4E-T antibodies (Bottom panel) followed by confocal microscopy. The nuclei were stained with DAPI. **C.** 293T cells treated with either vehicle or NaAsO_2_ as in (A) were stained with anti-LSM14A and anti-G3BP1 antibodies, followed by confocal microscopy. White arrows indicate PBs juxtaposed to SGs.

LSM14A has also been reported to localize to SGs in certain contexts^32^. To test this in 293T cells, we induced stress granule formation with sodium arsenite and performed co-staining for LSM14A and the SG marker G3BP1. No localization of LSM14A in SGs was observed under these conditions. Notably, PBs and SGs were frequently juxtaposed (Fig. 2C), consistent with prior observations that these dynamic assemblies engage in crosstalk and exchange components^33,34^.

To determine whether PBs are required for LSM14A-mediated innate immune enhancement, we first disrupted PB formation by knocking out DDX6 or 4E-T in 293T cells. As expected, loss of either factor abolished PB formation (Fig. 3A-D). Strikingly, however, LSM14A overexpression in DDX6- or 4E-T-deficient cells still robustly enhanced SeV-induced innate immune signaling, as measured by ISRE reporter activity, to levels comparable to wild-type cells (Fig. 3E and 3F). These findings indicate that intact PBs are not required for LSM14A’s immune-enhancing function.

**Fig. 3.**
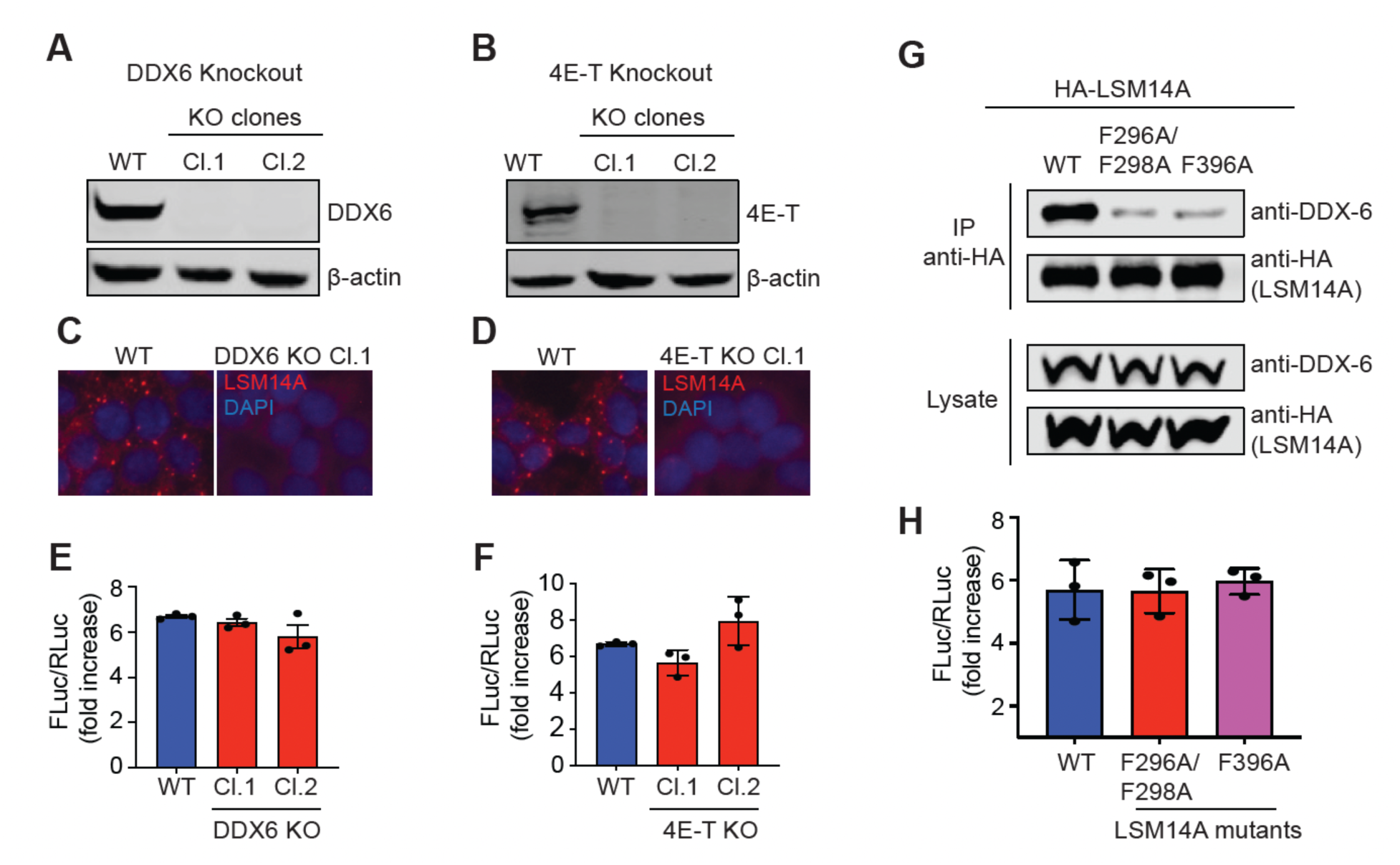
PB localization is not essential for LSM14A innate immune functions. **A, B.** 293T cell clones lacking DDX6 (A) or 4E-T (B) as demonstrated by western blot. **C, D.** Wild-type (WT) or one of the two knockout clones for DDX6 (C) or 4E-T for stained with anti-LSM14A antibodies to monitor the presence of PBs. DAPI was used to stain the nuclei. **E, F.** WT and knockout cells for DDX6 (E) or 4E-T (F) were co-transfected with pISRE-FLuc and pRL-RLuc, along with either empty plasmid or LSM14A, and the next day, were infected with SeV. The cells were harvested and subjected to the dual luciferase assay at 24 h post-infection. The FLuc:RLuc ratio from three biological replicates of LSM14A-transfected cells is presented as a fold increase over cells transfected with empty plasmid. The experiment was repeated twice. **G.** 293T cells overexpressing wild-type HA-LSM14A, HA-LSM14A F296A/F298A, or HA-LSM14A F396A were subjected to immunoprecipitation with anti-HA antibodies, followed by western blot analysis of DDX6. **H.** Dual luciferase assay of 293T cells transfected with either wild-type LSM14A or indicated mutants. The assay was performed essentially as in E and F. The experiment was repeated twice.

We next took a complementary approach by engineering LSM14A mutants defective in PB localization. Because LSM14A is recruited to PBs through interactions with DDX6 and 4E-T^27^, we targeted its C-terminal domain—implicated in DDX6 binding—and generated two mutants: one containing a combination of F296A and F298A substitutions and the other containing the single F396A substitution. These mutations abolished DDX6 interaction (Fig. 3G). However, the mutants retained full capacity to active the ISRE reporter upon overexpression, matching wild-type LSM14A (Fig. 3H). Collectively, these data reinforce the conclusion that PB localization is dispensable for LSM14A-driven innate immune enhancement.

LSM14A has been proposed to translocate to peroxisomes upon viral infection^10^; however, whether this relocalization is functionally linked to its innate immune function has remained unclear. To address this, we generated 293T cells lacking PEX19, a protein essential for peroxisome biogenesis^35^ (Fig. 4A). High-resolution stimulated emission depletion (STED) microscopy confirmed the absence of peroxisomes in PEX19 knockout cells (Fig. 4B). Strikingly, LSM14A overexpression in these cells induced ISRE activation to levels comparable to wild-type cells (Fig. 4C), indicating that peroxisomal localization is not required for LSM14A-mediated innate immune amplification.

**Fig. 4.**
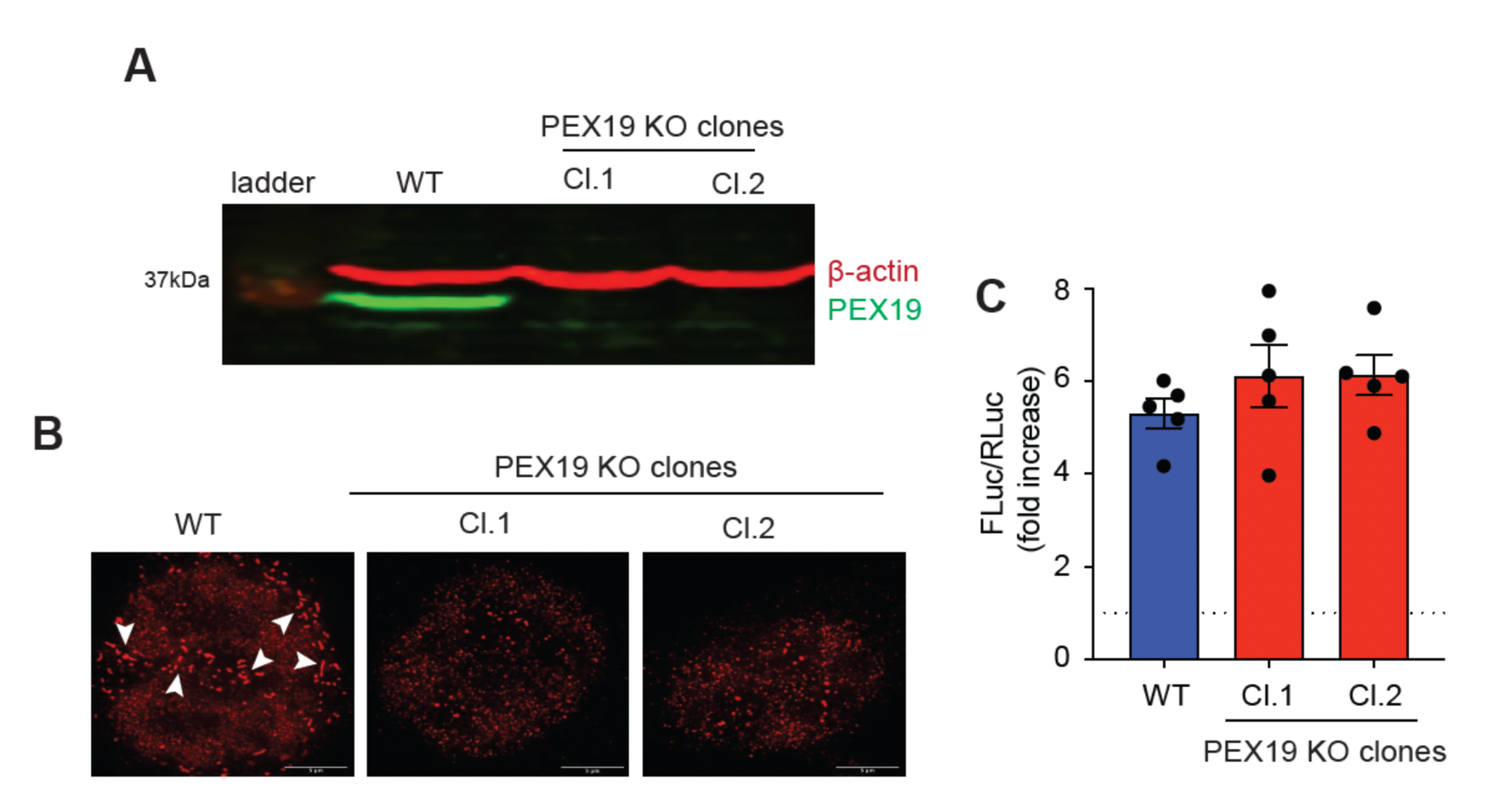
Peroxisome localization is not essential for LSM14A innate immune functions. **A.** 293T cell clones lacking PEX19 as demonstrated by western blot. β-actin was used as a loading control. **B.** Wild-type and PEX19-knockout 293T cells were stained with the peroxisome marker PMP70, and the cells were visualized by stimulated emission depletion (STED) microscopy. Since STED does not work best with DAPI, it was excluded from the staining. **C.** The ISRE activation experiment was performed as in Fig. 3E. The experiment was performed in quadruplicate.

### Minimal LSM14A protein length required for its innate immune functions

LSM14A consist of an N-terminal LSM domain, followed by an extended intrinsically disordered region (IDR; amino acids 84-290), and a largely unstructured C-terminal region harboring short motifs that mediate protein-protein and protein-RNA interactions required for PB assembly (Fig. 5A). Notably, LSM14A isoform 3, which differs from isoform 1 by a 40-amino acid deletion within the IDR (residues 139-179) (Fig. 5A), retained full capacity to activate ISRE, matching the activity of isoform 1 (Fig. 5B). This observation prompted us to directly test whether the IDR is required for LSM14A’s innate immune function.

**Fig. 5.**
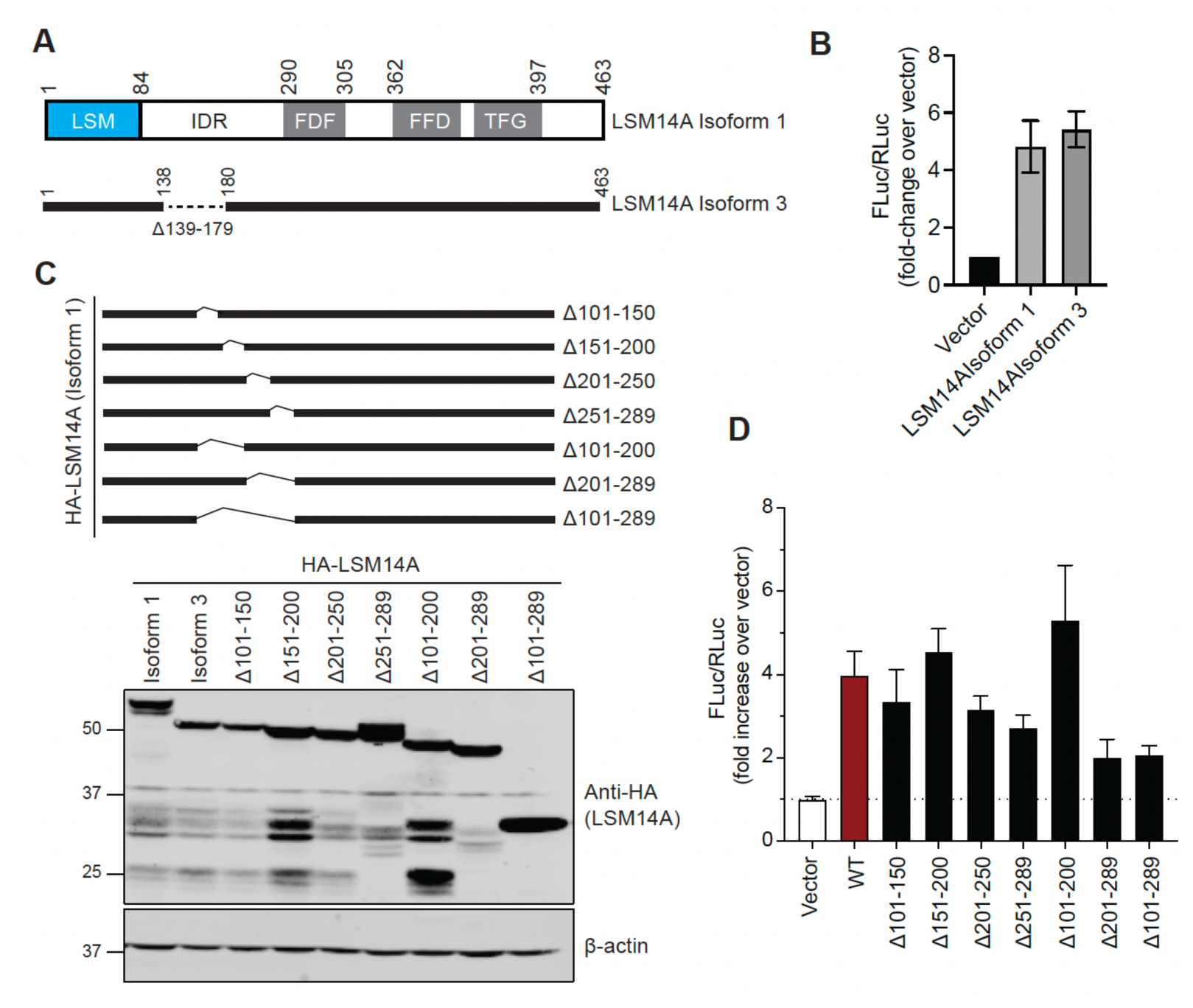
Determining the minimum protein length of LSM14A required for its innate immune function. **A.** Schematics of the LSM14A protein. Isoform 3 lacking amino acids 139-179 is also shown. **B.** ISRE activation by isoform 1 versus isoform 3 in 293T cells. The experiment was performed essentially as in Fig. 3E. **C.** Schematics of HA-LSM14A deletion mutants (top panel). 293T cells were transfected with indicated LSM14A mutants, and their expression was assessed by western blot with anti-HA antibodies at 48 hours post-transfection (bottom panel). **D.** ISRE activation by full-length LSM14A versus LSM14A deletion mutants. The bars represent mean ± SD of three biological replicates. The data in B and D are presented as fold-change compared to cells transfected with the empty vector.

To address this, we generated a series of deletion mutants lacking 50-100 amino acid segments across the IDR and assessed their ability to activate ISRE. All mutants were robustly expressed in 293T cells (Fig. 5C). Strikingly, the LSM14A Δ101-200 mutant—lacking most of the IDR—retained full activity, enhancing SeV-induced innate immune signaling to levels comparable to wild-type LSM14A. These findings indicate that large portions of the IDR can be removed without disrupting LSM14A function. Given the established role of IDRs in PB assembly^36^, these results further strengthen our conclusion that PB localization is not required for LSM14A’s innate immune activity.

### Interactomic analyses of LSM14A in SeV-infected cells

To define the PB- and peroxisome-excluded LSM14A interactome during SeV infection, we generated PEX19^-/-^ 293T cells stably expressing the Flag-tagged LSM14A Δ101-200 mutant under a doxycycline-inducible promoter. This mutant was specifically chosen to map LSM14A interactions outside of PBs and to minimize nonspecific interactions commonly mediated by intrinsically disordered regions. In parallel, we established control PEX19-deficient 293T cells expressing Flag-mKate under identical conditions. Following doxycycline induction, cells were either left uninfected or infected with SeV for 16 hours, after which Flag-tagged complexes were isolated and subjected to liquid chormatography followed by tandem mass spectrometry (LC-MS/MS). Principal Component Analysis (PCA) of the resulting proteomic datasets revealed clear separation between mKate and LSM14A interaction profiles under both mock and infected conditions (Fig. 6A), demonstrating that LSM14A engages in highly specific protein interactions. Consistent with this, bioinformatic analysis showed a markedly greater enrichment of proteins associated with LSM14A compared to mKate across conditions (Fig. 6B). Some of these proteins have previously shown to interact with LSM14A.

**Fig. 6.**
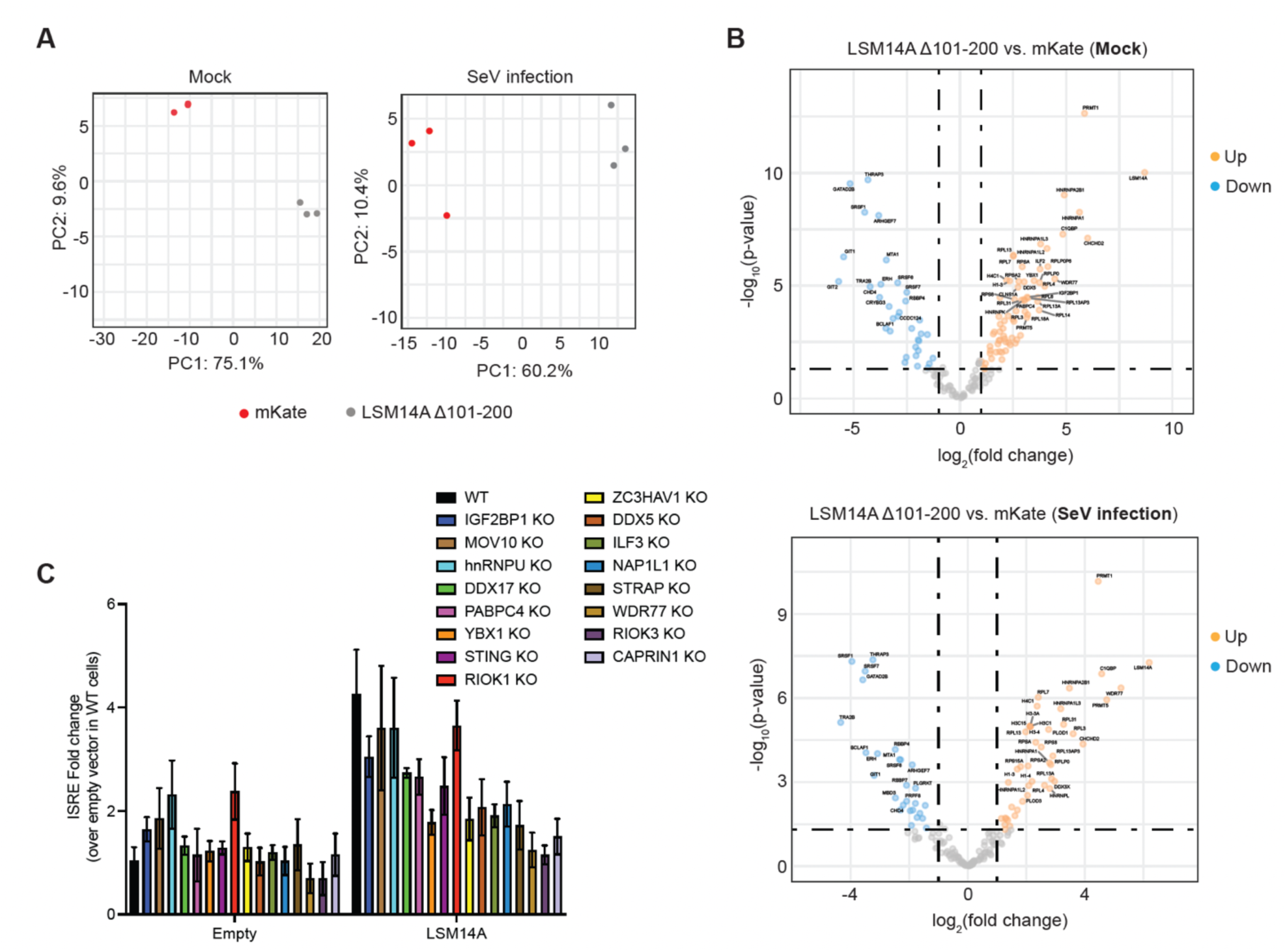
LSM14A interaction partners. **A.** PCA analysis. **B.** Volcano plot of differentially enriched proteins between Flag-LSM14A Δ101-200 and Flag-mKate in mock (upper panel) and SeV-infected 293T cells (bottom panel). Log_2_ fold change of Flag-LSM14A Δ101-200 over Flag-mKate are along the x-axis and - log_10_ *p*-values are plotted along the y-axis. **C.** Indicated genes were knocked out in 293T cells at the population level using the CRISPR/Cas9 system. The cells were then either transfected with an empty plasmid or LSM14A-expressing plasmid, infected with SeV, and subjected to the dual luciferase assay as described in Fig. 3E. The bars represent mean ± SD of four biological replicates.

To assess the functional relevance of these interactions, we prioritized candidate proteins, with a particular focus on those previously implicated in innate immune signaling, and systematically disrupted them in 293T cells using CRISPR/Cas9. We then evaluated LSM14A-dependent amplification of SeV-induced ISRE activation in these cells. Interestingly, loss of several candidate proteins abolished LSM14A’s ISRE-enhacing activity (Fig. 6C), indicating that these interactions are functionally critical for its innate immune function. Together, these findings define a specific LSM14A-centered interaction network and identify key host factors that are essential for its innate immune-enhancing activity, providing a foundation for future mechanistic investigations.

## DISCUSSION

LSM14A is a PB-associated, RNA-binding protein that regulates RNA metabolism. Li et al., reported that LSM14A can sense viral RNA, leading to the proposal that it functions in early innate immune detection, particularly at stages when canonical sensors such as RIG-I and MDA-5 are expressed at low levels^10^. This model suggests that incoming viral RNA may be trafficked to PBs, where LSM14A initiates early interferon (IFN) response. Our findings challenge this view. We demonstrate that PBs are dispensable for LSM14A-mediated amplification of innate immune signaling in 293T cells. More importantly, we found that LSM14A overexpression robustly potentiates innate immune activation not only in response to SeV infection or poly(I:C) stimulation, but also upon low-level expression of MAVS—which alone is insufficient to trigger signaling (data not shown). These results indicate that LSM14A can amplify innate immune responses even in the absence of exogenous RNA stimuli.

These observations however do not preclude a role for LSM14A’s RNA binding activity. Recent work shows that MAVS associates with host RNA to prime its signaling capacity^37^. By analogy, LSM14A may similarly depend on RNA binding, potentially outside of PBs. It is possible that LSM14A actively remodel host RNA to generate immunostimulatory species. Given its reported role in recruiting the mRNA decapping complex to the 5′ cap^27^, it is plausible that LSM14A overexpression enhances mRNA decapping, leading to the accumulation of uncapped RNAs that prime innate immune signaling and sensitize cells to external stimulie. Further experiments will be required to determine the extent to which LSM14A’s RNA binding activity underlies its antiviral function.

We would also like to note that depletion of DDX6 and 4E-T does not definitively rule out the possibility of LSM14A localized within mRNP assemblies. Prior work has shown that DDX6 depletion promotes SG formation^32^, and because LSM14A can localize to SGs^32^, it is plausible that, in DDX6-deficient cells, overexpressed LSM14A is redirected to SGs where it performs its innate immune function. Alternatively, although DDX6 or 4E-T depletion disrupts steady state PB formation (Fig. 3A), LSM14A overexpression may drive the assembly of non-canonical, PB-like compartments that provide a permissive environment for its function. This possibility is supported by observations that DDX6-deficient cells exposed to sodium arsenite, form PB-like assemblies containing key components, such as 4E-T, EDC4, and DCP1A^32^. Detailed analyses revealed that, when DDX6 is depleted, EDC4 and 4E-T segregate into distinct subassemblies with limited colocalization, suggesting that loss of DDX6-4E-T interactions alters the architecture of these structures. Similar remodeling is expected in 4E-T knockout cells. Consistent with this concept, studies in yeast have shown that PBs arise from a network of multivalent interactions, such that loss of individual components results in smaller, heterogenous assemblies, some of which may fall below the resolution of conventional microscopy^38^.

Our prior work demonstrated that depletion of MAVS abolishes LSM14A-mediated amplification of innate immune signaling following SeV infection, whereas LSM14A depletion does not impair MAVS-driven immune activation^16^. These findings place LSM14A upstream of, or at the level of, MAVS. Based on the observation that LSM14A relocalizes to the peroxisomes upon SeV infection^10^, it has been proposed that it enhances signaling by engaging peroxisome-associated MAVS. However, our data demonstrate that peroxisomes are dispensable for LSM14A function. It is possible that in peroxisome-depleted cells, LSM14A interfaces with MAVS at alternative subcellular sites, such as mitochondrial membranes, or through transient or indirect interactions. Notably, we did not detect enrichment of MAVS in LSM14A immunoprecipitates, under either basal or SeV-infected conditions. Similarly, LSM14A was not enriched in the MAVS pull-downs^37^, suggesting that any functional coupling is likely indirect or highly dynamic.

Finally, our proteomic analyses have uncovered a network of LSM14A-associated proteins that localize outside of peroxisomes and canonical PBs. Although their interactions and mechanisms of action remain to be fully defined, preliminary genetic perturbation studies indicate that several of these proteins are required for LSM14A-mediated immune amplification. Collectively, these findings support a model in which LSM14A operates within a flexible and context-dependent interaction network, independent of canonical compartments, to potentiate antiviral innate immunity.

## MATERIALS AND METHODS

### Cells and viruses

Human epithelial kidney HEK293T cells (ATCC CRL-3216) and human colorectal adenocarcinoma Caco-2 cells (ATCC HTB-37) were cultured in Dulbecco’s Minimum Essential Medium (DMEM) (Gibco; #11-995-065) supplemented with FBS (10%), penicillin (100 U/mL), and streptomycin (100 µg/mL) (Gibco; #10378016). Human bronchial epithelial BEAS-2B (ATCC CRL-3588) were cultured in Ham’s F-12K medium (Gibco; #11765054) supplemented with FBS (10%), penicillin (100 U/mL), and streptomycin (100 µg/mL). All cell lines were cultured at 37 °C with 5 % CO_2_ and were regularly verified to be free of mycoplasma.

The Sendai virus Cantell strain was purchased from Charles River Laboratories. Herpes simplex virus 1 was a kind gift from Dr. David Knipe (Harvard Medical School). The recombinant poliovirus Mahoney strain containing a catalytically inactive mutation in the 2A protease was produced by reverse genetics, using the strategy Schipper JG *et al.* employed to generate a CVB3 2A^pro^ mutant^29^. Briefly, the active site cysteine (C110) was substituted by alanine, and the 2A^pro^ cleavage site between VP1 and 2A was replaced with the 3CD cleavage site targeted by the 3C^pro^. Human seasonal coronavirus 229E containing enhanced green fluorescent protein (eGFP) as a replacement of the viral ORF4 gene was rescued using circular polymerase extension reaction (CPER).

### Antibodies and reagents

The antibodies used in this study include rabbit anti-LSM14A (Bethyl Laboratories; #A305-103A), mouse anti-LSM14A (Santa Cruz Biotechnology; #sc-398633), anti-DDX6 (Bethyl Laboratories; #A300-461A), anti-4E-T (Bethyl Laboratories; #A300-706A), anti-PEX19 (Proteintech; #14713-1-AP), anti-G3BP1 (Santa Cruz Biotechnology; #sc-365338), anti-PMP70 (Sigma-Aldrich; #SAB4200181), anti-HA (ABclonal; #AE008), anti-GAPDH (GeneTex; #GTX627408), and anti-β-actin (Thermo Fisher Scientific; #AM4302). The secondary antibodies used for western blot include IRDye 680RD goat anti-mouse (LI-COR; #926-68070) and IRDye 800CW goat anti-rabbit (LI-COR; #926-32211) and those used for IF include goat anti-rabbit Alexa Fluor 488 (Thermo Fisher Scientific; #A-11008), goat anti-rabbit Alexa Fluor 568 (Thermo Fisher Scientific; #A-11011), goat anti-mouse Alexa Fluor 488 (Themo Fisher Scientific; #A-11029), and goat anti-mouse Alexa Fluor 568 (Thermo Fisher Scientific; #A-11004).

Avicel was purchased from Dupont (#RC-581-NFDR080I), DAPI from Thermo Fisher Scientific (#D1306), Dual-Luciferase Reporter Assay System from Promega (#E1980).

### Plasmids

Transient expression plasmid (pcDNA3.1) and lentiviral vector (pLOC) expressing full-length LSM14A have been previously described^16^. These plasmids were used as templates to introduce point mutations or amino acid deletions into LSM14A. For doxycycline-inducible expression of LSM14A, the cDNA was cloned as a Flag-tagged construct into the enhanced PiggyBac (ePiggyBac) plasmid obtained from Dr. Ali Brivanlou (The Rockefeller University)^39^. Flag-tagged mKate2 was also cloned into ePiggyBac to be used as a control in LSM14A interactomic studies. The helper plasmid expressing the transposase cDNA was obtained from Dr. Ali Brivanlou (The Rockefeller University)^39^. MAVS cDNA was cloned into both pcDNA3.1 and pLOC using standard recombinant DNA technology.

The pISRE-FLuc plasmid, containing the Firefly luciferase reporter under an ISRE promoter, was from Stratagene (#219089) and pRL-RLuc, containing the CMV promoter-driven Renilla luciferase reporter, was from Dr. Stanley Lemon^40^. The small guide RNA (sgRNA) cloning plasmid pSpCas9(BB)-2A-Puro (pX459) was purchased from Addgene (#481239). The sgRNAs were designed using <u>crispor.tefor.net.</u> (Table 1) and cloned into pX459 plasmid for transfection into cells.

**Table 1:**
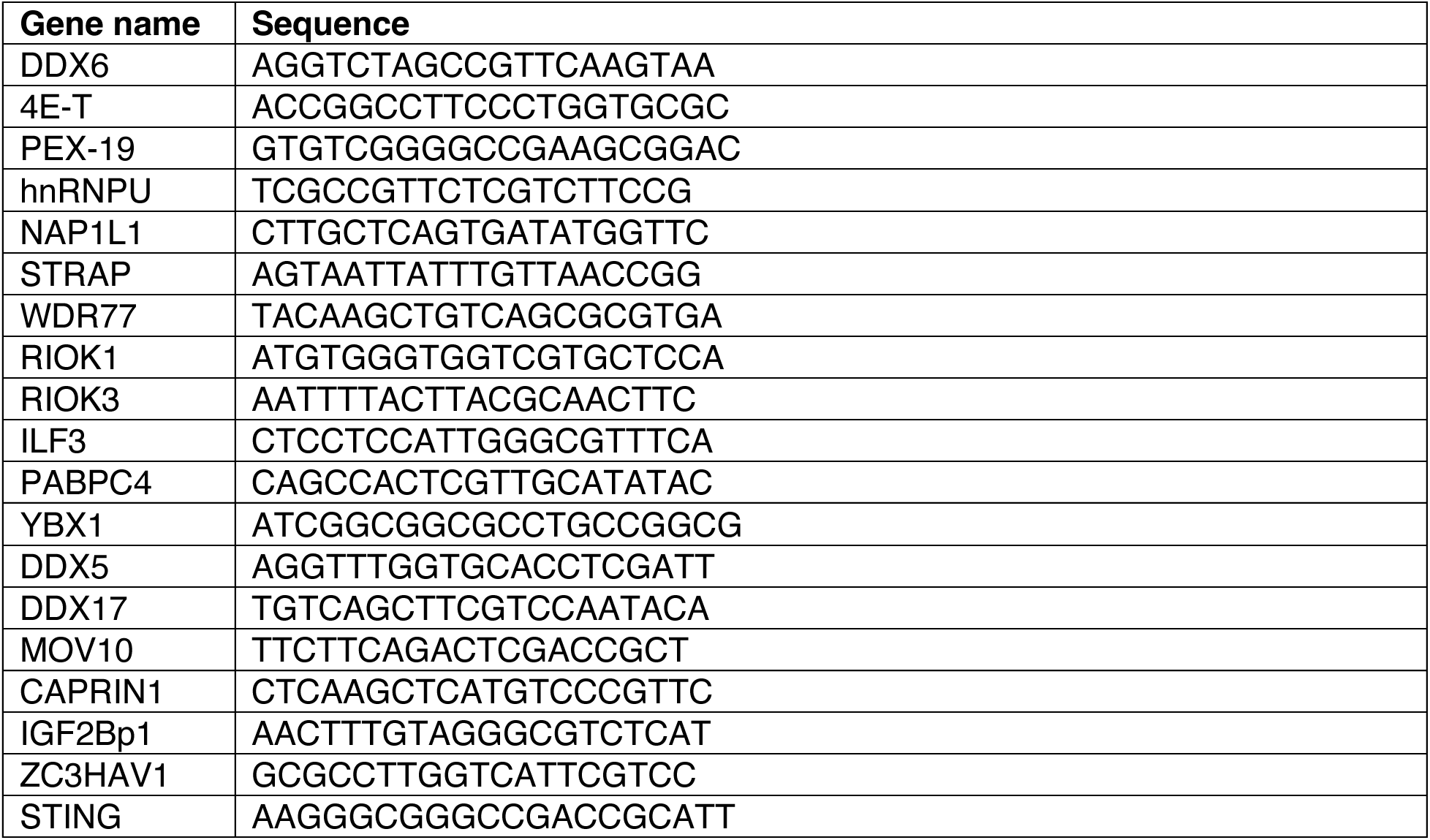
Nucleotide sequences of gRNAs used for genetic perturbations.

### Generation of knockout cells

We generated knockout cell clones essentially as previously described. Briefly, 293T cells were transfected with pX459 plasmid containing sgRNA using Lipofectamine 2000 DNA transfection reagent (Thermo Fisher Scientific; #11668030). We used two gRNAs per gene to increase the probability of complete knockout. Thirty-six hours after transfection, the cells were treated with 2.5 µg/mL puromycin for 48h and then recovered for three days in puromycin-free medium. The single cell clones were obtained by seeding the cells into 96-well plates at a density of 0.7 cells per well. The resulting cell clones were screened for gene expression by Western blot.

### Luciferase assay

Dual luciferase assay was performed to evaluate the ISRE activation. Cells were seeded into 96-well plates at a density of 40,000 cells per well (293T) or 25,000 cells per well (BEAS-2B), and the next day, were transfected with DNA using FuGENE HD (Promega; #E2311). Each well received 100 ng pISRE-FLuc, 2.5 ng pRL-RLuc, and 50 ng of either pcDNA3.1, pcDNA3.1-LSM14A, or pcDNA3.1-MAVS. Twenty-four hours later, the cells were either infected with SeV or left uninfected, and at 24 hours post-infection, were lysed in 50 µl of 1X passive lysis buffer (PLB) from the Dual Luciferase Assay Kit (Promega; #E1980). 10 µL of the lysate was transferred into a white 96-well plate (Corning) and the luciferase signal was measured using VarioSkan Lux Multimode Microplate Reader (Thermo Fisher Scientific).

### Measuring antiviral activity of LSM14A

Caco-2 and BEAS-2B cells were engineered to stably overexpress empty plasmid, LSM14A, or MAVS. Caco-2 cells were infected with PV-2A^mut^ and BEAS-2B cells with 229E and HSV-1. All infections were performed at a multiplicity of infection (MOI) of 0.01, and the virus adsorption was carried out for 1h at 37°C, following which the cell monolayers were washed twice with 1X PBS, and the fresh medium was added to each well. The culture medium was collected at 24 hours post-infection (hpi). PV-2A^mut^, 229E, and HSV-1 were titrated on HeLa, HuH-7, and Vero E6 cells, respectively.

### Immunofluorescence

For confocal microscopy, cells were seeded on poly-L-lysine-coated glass coverslips. Following incubation at 37°C for 24 hours, the cells were either left untreated or treated with 100 µM sodium arsenite for 1h at 37°C. For overexpression studies, the cells were transfected with 100 ng of plasmids using Lipofectamine 3000 (Thermo Fisher Scientific; #L3000015). They were fixed in 10% formaldehyde for 15 min at room temperature (RT) and permeabilized in 1X PBS containing 0.1% Triton X-100 for 30 min at RT. Blocking was performed with the blocking buffer containing 1X PBS, 0.1% Triton X-100, 1% BSA, and 2% goat serum for 1h at RT, followed by overnight incubation at 4°C with primary antibodies diluted in blocking buffer (mouse anti-LSM14A, 1:100; rabbit anti-LSM14A, 1:500; anti-DDX6, 1:100; anti-4E-T, 1:100; anti-G3BP1, 1:100). The cells were extensively washed in 1X PBS and incubated with secondary antibodies for 1h at RT. Nuclei were counterstained with DAPI for 10 min.

### STED Imaging

The 293T cells grown on poly-L-lysine-coated cover slips were fixed with 4% paraformaldehyde (PFA) for 15 min at RT and permeabilized for 15 min with 1X PBS + 0.1% Triton X-100 at RT. Cells were then blocked for 1 hour at RT using 1X PBS with 0.1% Triton X-100, 1% BSA, and 2% goat serum. Cells were incubated overnight at 4°C with anti-PMP70 antibody (1:250) diluted in blocking buffer, followed by 5X washing with 1X PBS. The cells were then incubated with goat anti-mouse IgG conjugated STAR RED (1:250; Abberior; #STRED-1001) for 1 hour at RT. Cells were again washed five times with 1x PBS. Coverslips were mounted using ProLong Gold Antifade Mountant (Invitrogen). STED imaging was acquired using an Abberior STED super-resolution microscope.

### Immunoprecipitation

293T cells stably overexpressing HA-LSM14A, either wild-type or mutants, were subjected to IP using anti-HA beads (Pierce Anti-HA Magnetic Beads; #88836). Cell expressing HA-only served as a control. Briefly, the cells were lysed in a buffer containing 50 mM Tris-HCl pH 7.4, 150 mM NaCl, 1 mM EDTA, 0.5% NP-40, protease inhibitors, and phosphatase inhibitors. The lysates were clarified to remove the debris and incubated with anti-HA magnetic beads for 2h at 4°C, followed by washing of beads with 1X PBS. The bound proteins were eluted in 90 µl of 1X LDS buffer by incubating at 70°C for 10 min and revealed by Western blot.

### LSM14A interactomic analysis by affinity purification followed by mass spectrometry (AP/MS)

293T cells engineered to express Flag-mKate and Flag-LSM14A Δ101-200 under a doxycycline-inducible promoter were seeded in 10 cm dishes (4 dishes per condition). The next day, there were treated with doxycycline (1 µg/mL), and 36h later, were stimulated or not with SeV. Twenty-four hours post-infection, the cells were rinsed with 10 ml of ice-cold 1X PBS and collected in 15-mL falcon tubes by scraping in 1 mL of ice-cold 1X PBS. The cells were pelleted by centrifugation at 300 xg for 5 min at 4°C and the pellet was homogenized in 1 ml of lysis buffer containing 50 mM Tris-HCl pH 7.4, 150 mM NaCl, 1 mM EDTA, 0.5% NP-40, protease inhibitors, and phosphatase inhibitors. The lysates were transferred to 1.5 mL Eppendorf tubes and incubated on dry ice for 15-20 min followed by quick thawing at 37°C for 5-10 min. This freeze-thaw cycle was repeated again, and the lysates were centrifuged at 13,000 xg for 15 min. The supernatants were incubated with anti-Flag magnetic beads (Pierce Anti-DYKDDDDK Magnetic Agarose; #A36797) on a rotator at 4°C for 2h. The beads were extensively washed (3X in 1 ml of lysis buffer) and transferred to new LoBind tubes followed by one additional wash. The beads were then incubated in 90 µl of 1X LDS at 70°C for 10 min with constant shaking (1000 rpm), the eluted proteins were transferred to new Eppendorf tubes, and 10 µl of 1M DTT was added to reduce proteins. Samples were stored at -80°C until downstream analysis.

### In-gel enzymatic digestion

Eluted proteins were separated on precast 4-12% SDS gels, followed by gel staining with Coomassie blue for 2 h. Gel lanes were sliced into 4 large bands, and each band was cut into smaller pieces and added to a 96-well plate containing 50 mM ammonium bicarbonate. The gels were destained using a 50% Acetonitrile/50% 50 mM ammonium bicarbonate solution 3-4 times until the gel pieces turned white, followed by dehydration using 100% acetonitrile. The gel pieces were then air dried and reduced with dithiothreitol (DTT) at 56°C for 30 min, alkylated with iodoacetamide (IAA) at RT for 30 min, and subjected to trypsin-LysC (Promega) digestion overnight at 37°C. The peptides were extracted from each gel band, and the solution was dried by vacuum centrifugation (SPD1010 Speedvac system, Thermo Savant). The dried samples were resuspended in 2% ACN/water/0.1% TFA and passed through C-18 ZipTips (Thermo Scientific); the cleaned peptides were eluted with 60% ACN/water/0.1% TFA and dried by vacuum centrifugation. The cleaned peptides were stored at -20 °C until analyzed by liquid chromatography-tandem mass spectrometry (LC-MS/MS).

### LC-MS/MS proteomic analysis

Nano-LC-MS/MS separation was performed using a NanoAcquity M-Class UPLC (Waters Technology) coupled to an Orbitrap Fusion Lumos MS (Thermo Fisher Scientific). Reversed-phase nanoEase V/M Symmetry C18 trap column with a 180 μm internal diameter and a nanoEase M/Z HSS T3 column with a 75 μm internal diameter were used with a 120-min method at a constant flow rate of 0.5 µL/min. The MS acquisition was performed over a mass range of m/z 350-2000, with a maximum injection time of 50 ms and a resolution of 120 K. The normalized automatic gain control (AGC) target was set to 100% with a value of 4e^5^, and dynamic exclusion was set to 10s. Tandem MS was performed using an isolation window of m/z 1.6, a microscan of 1, a resolution of 30 K, a maximum injection time of 60 ms, and an AGC target of 5e^5^. A fixed normalized collision energy (NCE) in the high collision energy dissociation (HCD) mode of 27 V was used. All samples were randomized, processed, and acquired on the instrument simultaneously to minimize bias. Routine external standards, including a Peptide Retention Time Calibration Mixture (Thermo Fisher Scientific) and a HeLa Protein Digest Standard (Thermo Fisher Scientific), were run alongside the samples.

### Proteomic data analysis

The raw LC-MS/MS data were analyzed using available commercial PEAKS Studio v12 software (Bioinformatics Solutions, Inc., Waterloo, ON, Canada) against the UniProt/SwissPro database for homo sapiens taxonomy with a precursor mass tolerance of 10 ppm and MS/MS of 0.02 Da. Trypsin/LysC was selected as the enzyme with up to two missed cleavages. Carbamidomethylation modification was selected as a fixed modification. Protein and peptide identification was performed at a 1% false discovery rate. The label-free quantification was achieved using PEAKS Studio Quantification- a label-free module with a setting of mass error tolerance of 20 ppm and an autodetected retention time shift tolerance with a feature intensity ≥ 1e6, and all abundances were normalized using total ion chromatograms (TICs) by the software.

### Bioinformatic analysis of the proteomic dataset

Differential enrichment analysis was performed in R using the DEP package (v.1.28.0)^41^, which builds on limma for protein-wise linear modelling with empirical Bayes moderation^42^. Raw label-free quantification (LFQ) intensities were filtered to remove decoys and contaminants, log_2_-transformed, normalized by variance-stabilizing transformation, and missing values were imputed using a left-shifted Gaussian. Prior to pairwise comparisons, outlier samples were excluded based on PCA clustering and LFQ intensity distributions. Pairwise contrasts were then generated by comparing LSM14A Δ101-200 against its matched mKate control under each treatment conditions (Mock or SeV).

To identify high-confidence interactors, the list of candidate genes was filtered using two complementary specificity scores. The first filtering was performed using Contaminant Repository for Affinity Purification (CRAPome)^43^, which is a curated database of background proteins compiled from hundreds of negative control AP-MS experiments. It assigns each protein a frequency-and-abundance score (FC-A) reflecting how often it co-purifies non-specifically across unrelated pulldowns. Candidates were retained only if their FC-A score exceeded 3.0. The second filtering was based on the MiST (Mass Spectrometry interaction STatistics) score. MiST integrates abundance, reproducibility, and specificity into a single score (0-1). To capture proteins specifically enriched for LSM14A Δ101-200, a candidate was required to score ≥ 0.6 in LSM14A Δ101-200 pull downs and ≤ 0.4 in matched mKate controls.

The final list consisted of proteins that satisfied the following conditions in at least one pair-matched contrast: raw p-value < 0.05, |log_2_FC| ≥ 1.5, CRAPome FC-A ≥ 3.0, MiST ≥ 0.6 in mut5, and MiST ≤ 0.4 in mKate. Visualizations generated with ggplot2 (v 4.0.2), ggrepel (v0.9.8), and pheatmap (v1.0.13); heatmap values represent log_2_-normalized intensities, with rows hierarchically clustered and samples grouped by condition.

### Statistical analysis

Statistical analyses were performed using GraphPad Prism.

## ACKNOWLEDGEMENTS

We thank Dr. Frank Kuppeveld (Utrecht University, The Netherlands) for providing the CVB3 infectious clone with a catalytically inactive mutation in 2A protease^29^. This work was supported by Boston University startup funds (to M.S.), and by the National Institute of Health grant R01 AI170877 (to M.S.).

## AUTHOR CONTRIBUTION

Conceptualization, M.S.; Methodology, A.W., H.H., M.S., D-Y.C., C-M.S., J.D.P., M.S., Z.D.; Validation, A.W., H.H., D-Y.C.; Mass Spectrometry, M.S.; Bioinformatic Analysis, H.H., Writing-Original Draft, M.S.; Writing-Review & Editing, all authors; Supervision, M.S., S.M.L.; Funding Acquisition, M.S.

